# Non-destructive Spatial Reconstruction of Plant Leaf Starch Using Reduced-Band SWIR Spectroscopy and Chemometric Modeling

**DOI:** 10.64898/2026.06.02.728709

**Authors:** Ahmed Glili, Sajid Bangash, Matthias König, Daniel Šmít, Jan Dräger, Hee Sung Kang, Berit Ebert, Alois Knoll, Malte C. Gather, Stefan Hey

## Abstract

Non-structural carbohydrates (NSCs) are central to plant carbon allocation and physiological regulation, yet their quantification typically relies on destructive biochemical assays that lack spatial resolution. Here, we developed a shortwave infrared (SWIR) hyperspectral imaging workflow for non-destructive estimation and spatial reconstruction of starch-associated variation in strawberry leaves. The workflow combined automated hyperspectral segmentation, spectral preprocessing, Partial Least Squares Regression (PLSR), and constrained wavelength selection. Sample-level spectra extracted from 114 strawberry leaf samples grown across three different metabolic conditions were paired with destructive starch measurements and used to train models across the 900–1750 nm spectral range. A constrained greedy band-selection strategy revealed that predictive performance approached a plateau at approximately 12 wavelengths, indicating substantial spectral redundancy within the full hyperspectral dataset. The final reduced-band model achieved a cross-validated coefficient of determination (R^2^) of 0.771 ± 0.066 and a root mean squared error (RMSE) of 0.743 ± 0.098 mg g^−1^ fresh weight using repeated stratified 5-fold cross-validation. Pixel-wise application of the final model generated spatial starch-associated maps that preserved pronounced intra-leaf heterogeneity, including vein-associated spatial structure. These results demonstrate that starch-associated spectral information can be reconstructed from a constrained reduced-band SWIR framework while retaining sufficient predictive performance for spatial mapping. The identified wavelength reduction supports the feasibility of deployable multispectral systems for non-destructive carbohydrate sensing in plant phenotyping applications.

## 2 Introduction

Non-structural carbohydrates (NSCs) are central intermediates in plant carbon allocation, linking photosynthetic carbon assimilation with growth, respiration, storage, and stress responses. In leaves, starch functions as a transient carbon reserve that accumulates during the photoperiod and is remobilized to sustain metabolism and growth when photosynthesis is limited. The regulation of leaf starch turnover is closely coordinated with diel carbon availability, and perturbations in starch synthesis or degradation can affect growth dynamics and plant fitness (Graf and Smith, 2011; Hartmann and Trumbore, 2016; Stitt and Zeeman, 2012; Sulpice *et al*., 2009; Thalmann and Santelia, 2017).

Despite its physiological importance, starch is commonly quantified using destructive biochemical assays (Landhäusser *et al*., 2018; Smith and Zeeman, 2006). These approaches provide accurate bulk measurements, but they are laborious, terminal, and poorly suited for repeated measurements of the same tissue over time. They also integrate heterogeneous anatomical regions into a single sample-level value, thereby obscuring spatial variation within leaves. This limits the ability to resolve how starch accumulation differs between veins, mesophyll regions, leaf margins, or local source–sink domains (Landhäusser et al., 2018; Smith and Zeeman, 2006). Non-destructive spectroscopic approaches have been explored as an alternative, including contact-probe reflectance measurements on excised leaf tissue (Frey *et al*., 2020), but these methods still rely on spatially averaged spectra and do not provide the within-leaf spatial resolution that imaging-based approaches can offer.

Optical spectroscopy offers a route toward non-destructive biochemical phenotyping. In particular, near-infrared (NIR) and shortwave-infrared (SWIR) reflectance contain information related to overtone and combination absorptions of O–H, C–H, and N–H bonds and have been widely used to estimate plant and food-quality traits. Hyperspectral imaging extends this principle spatially by combining spectral information with image-based tissue localization, allowing biochemical predictions to be mapped across intact samples rather than averaged across entire organs. Prior work has demonstrated the feasibility of hyperspectral approaches for estimating plant biochemical traits, including starch content in leaves and storage organs (Curran, 1989; Frey *et al*., 2020; Golic *et al*., 2003; Kokaly *et al*., 2009; Ramirez *et al*., 2015; Sarić *et al*., 2022).

Despite demonstrated feasibility, full hyperspectral systems remain difficult to translate into operational phenotyping or greenhouse monitoring platforms. They generate large data volumes, require careful calibration and preprocessing, and are typically more expensive and complex than multispectral systems. For practical deployment, it is therefore important to determine whether the relevant biochemical information is distributed broadly across the full spectrum or concentrated within a limited number of informative wavelength regions. Reduced-band models can preserve much of the predictive information while enabling simpler optical designs, faster image acquisition, and lower-cost instrumentation (Bai *et al*., 2024; Burnett *et al*., 2021; Liu *et al*., 2020; Mehmood *et al*., 2012; Sarić *et al*., 2022).

A further challenge is that spectral models are often developed from sample-level mean spectra, whereas the biological motivation for imaging is spatially resolved interpretation: starch accumulation in leaves is not anatomically uniform, with gradients observed between mesophyll cells, bundle sheath cells, and vascular tissue, and differences between adaxial and abaxial surfaces linked to light exposure (Tsai *et al*., 2009). This creates a methodological gap between bulk destructive reference measurements and pixel-wise prediction maps. Spatial mapping from models calibrated on bulk tissue therefore requires careful segmentation, preprocessing, and interpretation, particularly when anatomical heterogeneity and mixed pixels at tissue boundaries may influence local reflectance (Burnett *et al*., 2021; Frey *et al*., 2020; Pandey *et al*., 2017; Zhang *et al*., 2023).

Here, we developed a SWIR hyperspectral imaging workflow for non-destructive estimation and spatial reconstruction of leaf starch content. The workflow combines hyperspectral preprocessing, spectral-feature-based leaf segmentation, sample-level PLSR calibration against destructive starch measurements, constrained wavelength selection, and pixel-wise application of the final reduced-band model. We hypothesized that starch-associated spectral information is concentrated within a limited number of SWIR regions and that a constrained reduced-band model can retain predictive performance while enabling spatial visualization of starch-associated variation within intact leaves. Indeed, our results support that starch-associated spectral information is largely concentrated within a limited number of SWIR regions, with predictive performance approaching a plateau at approximately 12 selected wavelengths, and that a constrained reduced-band model retains sufficient predictive capacity for spatial reconstruction of starch-associated variation within intact strawberry leaves.

## 3 Results

### 3.1 Hyperspectral Preprocessing, Segmentation, and Spectral Structure

To capture a broad physiological range of starch concentrations, strawberry leaves were sampled across three source–sink treatments and across 11 time blocks spanning a 16 h photoperiod, yielding 114 matched imaging–biochemistry samples covering both diurnal NSC dynamics and treatment-driven variation. Each leaf was imaged within one minute of detachment with a push-broom SWIR hyperspectral camera (900–1750 nm) under broadband halogen line illumination (Fig. 1 A, B) and subsequently analyzed enzymatically for total starch to provide reference values for model calibration. This dataset formed the basis for all downstream analyses. We first developed a preprocessing and segmentation workflow to enable extraction of leaf tissue from calibrated hyperspectral image cubes across all evaluated samples and imaging conditions. Spatial and spectral preprocessing were applied to reduce sensor noise and reflectance artifacts while preserving physiologically relevant spectral variation within the leaf tissue. Subsequent multi-feature segmentation combined complementary spectral descriptors with voting-based thresholding to generate robust binary leaf masks for downstream spectral analysis (Fig. 1A).

**Figure 1.**
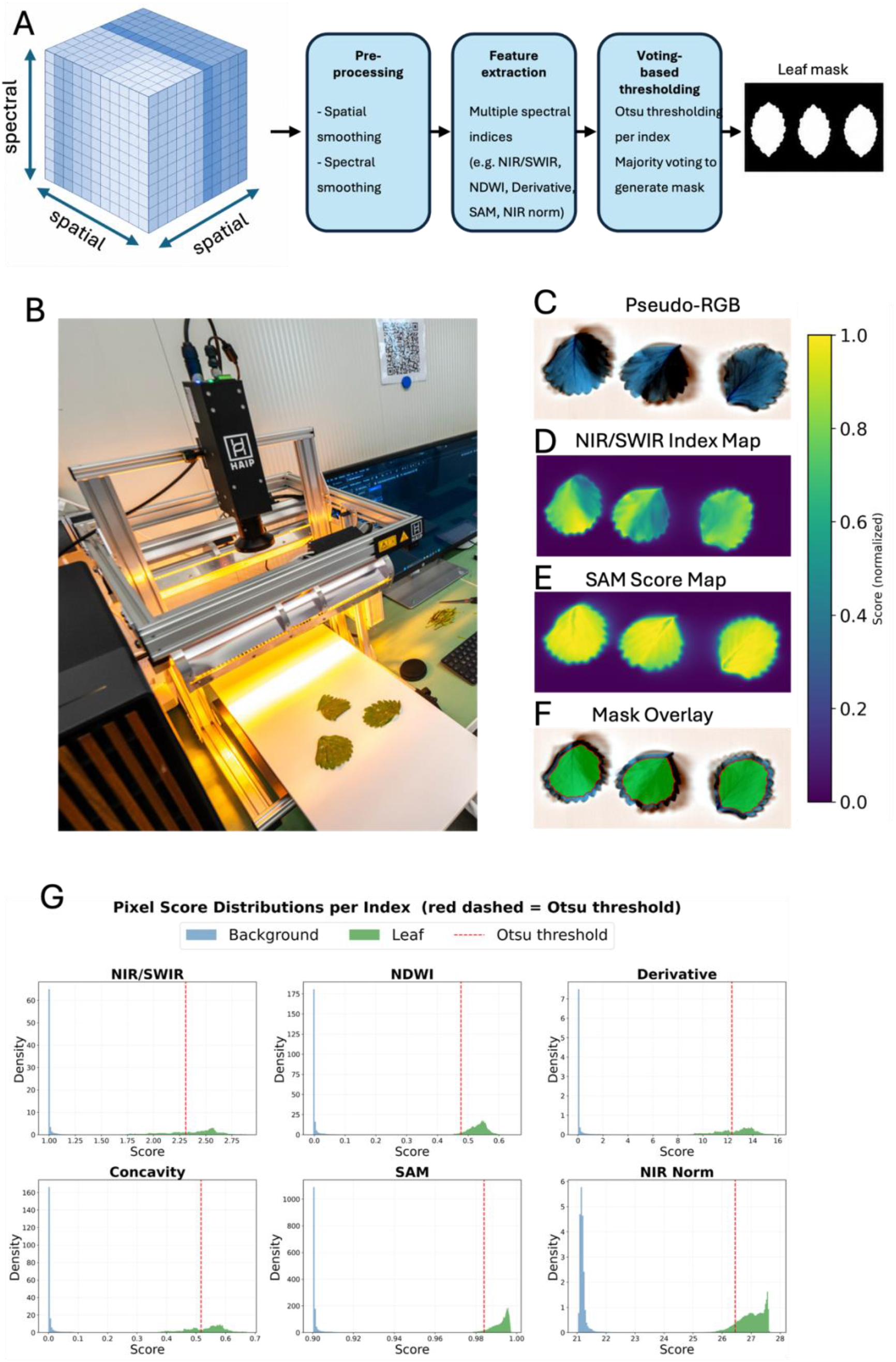
Hyperspectral imaging setup and preprocessing and segmentation workflow for leaf tissue extraction. (A) Overview of the preprocessing and segmentation pipeline applied to calibrated hyperspectral image cubes. The workflow consisted of spatial and spectral preprocessing, computation of multiple spectral descriptors, and voting-based thresholding for binary mask generation. Preprocessing included spatial smoothing and spectral smoothing. Feature extraction incorporated complementary spectral indices, including NIR/SWIR ratios, NDWI, spectral derivatives, Spectral Angle Mapper (SAM) similarity metrics, and normalized NIR deviation features. Otsu thresholding was applied independently to each descriptor map, followed by majority-vote aggregation to generate final leaf masks. (B) Photograph of the SWIR hyperspectral imaging setup, showing the line-scan camera mounted above a translating sample stage with halogen line illumination and example strawberry leaflets in the imaging field. (C) Pseudo-RGB representation of example strawberry leaflets reconstructed from the hyperspectral cube. (D) Representative NIR/SWIR index map highlighting strong spectral separation between leaf tissue and background regions. (E) SAM-based similarity map showing homogeneous spectral clustering within leaf interiors. (F) Final segmentation mask overlay demonstrating accurate preservation of leaf morphology and boundary structure. The shared colorbar in panels D–F displays normalized descriptor scores. (G) Pixel score distributions for the six evaluated spectral descriptors (NIR/SWIR, NDWI, Derivative, Concavity, SAM, and NIR Norm). Blue and green histograms represent background and leaf pixels, respectively, while red dashed lines indicate Otsu-derived threshold values. Clear bimodal separation was observed across all evaluated feature domains, supporting robust automated segmentation without manual parameter tuning.

Strong spectral discrimination between leaf and background pixels was observed across all evaluated feature domains (Fig. 1G). Pixel score distributions exhibited clear bimodal separation, enabling automated thresholding without manual parameter tuning. Among the evaluated descriptors, Spectral Angle Mapper (SAM)-based similarity metrics produced particularly distinct clustering behavior, with leaf pixels concentrated near unity and background pixels occupying substantially lower score ranges. Similar separation patterns were observed for derivative-based metrics and normalized NIR deviation features, whereas water-sensitive indices such as Normalized Difference Water Index (NDWI) additionally captured the strong SWIR absorption behavior associated with hydrated plant tissue (Fig. 1G).

The Otsu-derived thresholds consistently localized within low-density transition regions between foreground and background classes, indicating favorable class separability and supporting the robustness of the automated segmentation procedure. Spatial feature maps closely corresponded to visible leaf morphology, with elevated responses restricted primarily to leaf tissue while background regions remained near baseline values (Fig. 1D-F). Before further analysis, the generated masks were eroded, i.e. reduced in size, to exclude the region close to leaf boundaries which effectively prevented contamination of the data by local transitions in reflectance and non-uniform illumination conditions (Fig. 1F; Otsu, 1979).

### 3.2 Modeling Workflow and Constrained Band Selection

Sample-level spectra were extracted from the eroded leaf masks and paired with corresponding destructive starch measurements to construct the calibration dataset (Fig. 2). Prior to modeling, spectra were normalized using Standard Normal Variate (SNV) transformation to reduce multiplicative scattering effects and baseline variability associated with leaf morphology and illumination differences. Partial Least Squares Regression (PLSR) was used to model the relationship between hyperspectral reflectance and starch concentration. Model dimensionality was optimized using repeated stratified 5-fold cross-validation, in which samples were grouped into low-, medium-, and high-starch bins to preserve the response distribution across folds. Cross-validation performance stabilized at seven latent variables, which were subsequently retained for all downstream analyses. To identify a reduced spectral subset suitable for multispectral implementation, an initial full-spectrum PLSR model was trained using all available wavelength bands. Wavelengths were then ranked according to the absolute magnitude of their regression coefficients. A constrained greedy selection procedure was subsequently applied to iteratively select informative wavelengths while maintaining spectral diversity across the measured SWIR range. Inspection of the full-spectrum PLSR coefficients showed that the most influential wavelengths clustered within narrow contiguous regions, which would have caused an unconstrained greedy selector to draw an excessive number of adjacent bands from the same region. To avoid this redundancy and ensure that the reduced subset retained information from across the measured SWIR range, the selection procedure enforced both regional coverage and minimum spectral spacing constraints. Specifically, at least one wavelength was required from each predefined spectral region (900–1100 nm, 1100–1450 nm, and 1450–1748 nm), while neighboring selected wavelengths were separated by a minimum distance of 40 nm. If the target number of bands could not be reached under the initial spacing criterion, the minimum separation threshold was progressively relaxed in 5 nm increments. (Fig. 2; Barnes *et al*., 1989; Burnett *et al*., 2021; Geladi and Kowalski, 1986; Wold *et al*., 2001).

**Figure 2.**
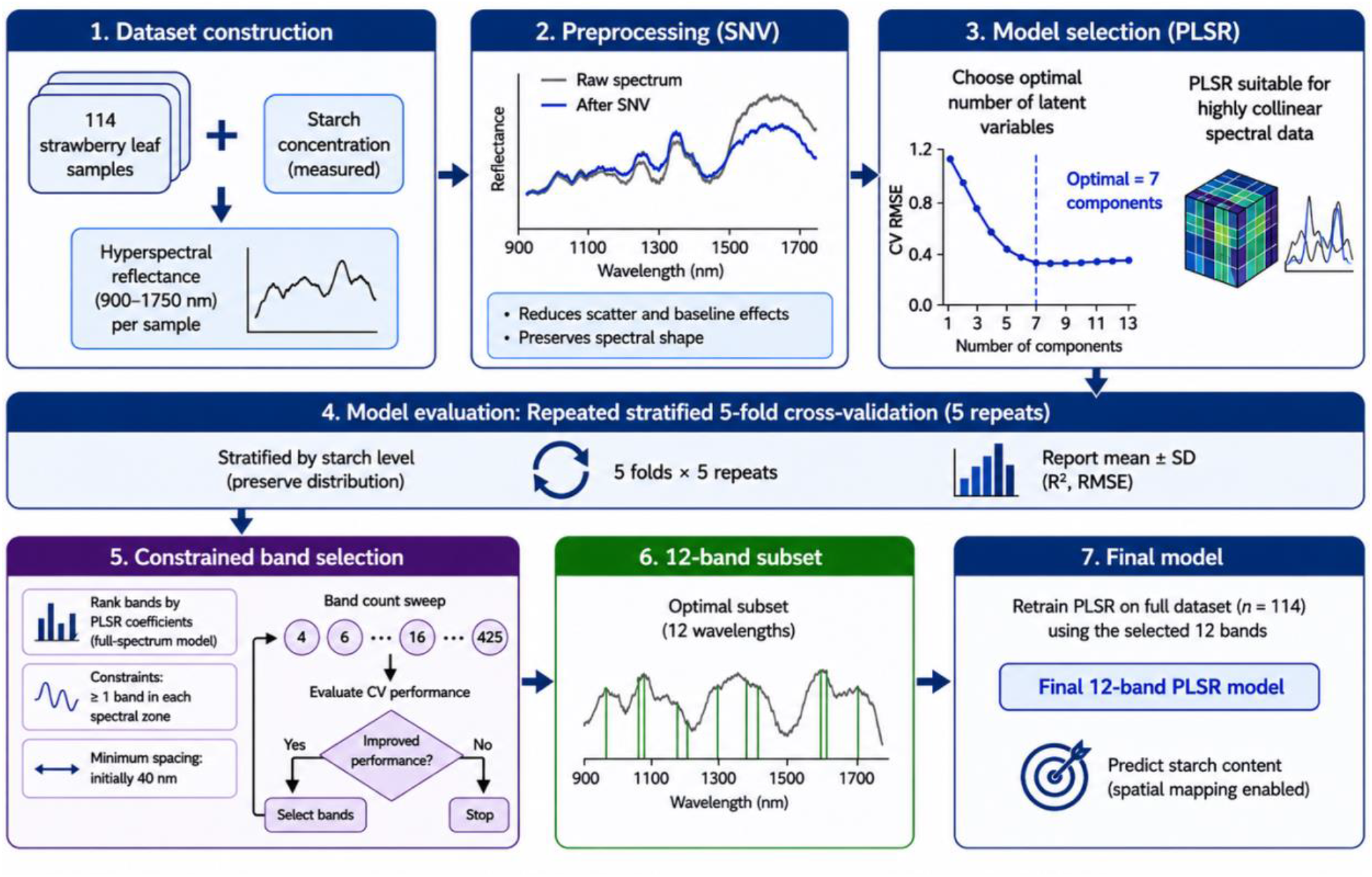
Modeling workflow for reduced-band starch prediction from SWIR hyperspectral data. (1) Dataset construction from 114 strawberry leaf samples. For each sample, hyperspectral reflectance spectra (900–1750 nm) were paired with destructive starch concentration measurements to generate the calibration dataset. (2) Spectral preprocessing using Standard Normal Variate (SNV) normalization to reduce multiplicative scattering effects and baseline variability while preserving spectral shape information. (3) Partial Least Squares Regression (PLSR) model selection and optimization. The optimal number of latent variables was determined by minimizing cross-validated root mean squared error (RMSE), resulting in a final model with seven components. (4) Model evaluation using repeated stratified 5-fold cross-validation (5 repeats). Samples were stratified according to starch concentration to preserve the response distribution across folds. Mean and standard deviation of cross-validated R^2^ and RMSE were recorded. (5) Constrained band-selection workflow. Wavelengths were ranked according to the absolute magnitude of PLSR regression coefficients obtained from the full-spectrum model. Candidate band subsets were generated under two constraints: (i) at least one wavelength per predefined spectral region and (ii) a minimum spectral spacing of 40 nm between selected wavelengths. Multiple candidate band counts were evaluated iteratively using cross-validation performance. (6) Final reduced-band configuration consisting of 12 selected wavelengths spanning multiple SWIR regions associated with starch-related spectral structure. (7) Retraining of the final reduced-band PLSR model on the complete dataset using the selected wavelengths, enabling downstream prediction and spatial starch reconstruction from hyperspectral image data.

### 3.3 Reduced-Band Selection and Spectral Contribution Analysis

Analysis of the reduced-band model revealed that several selected wavelengths localized within spectral regions exhibiting elevated latent loading structure and increased Variable Importance in Projection (VIP) scores (Fig. 3A). Major contributing regions were concentrated near approximately 930 nm, 1200 nm, and longer SWIR wavelengths above 1500 nm. Regression coefficient analysis further demonstrated that predictive information was distributed across multiple spectral neighborhoods rather than confined to isolated wavelength peaks (Fig. 3D). Together, these observations indicate that starch-associated prediction emerged from multivariate spectral covariance patterns spanning several SWIR domains.

**Figure 3.**
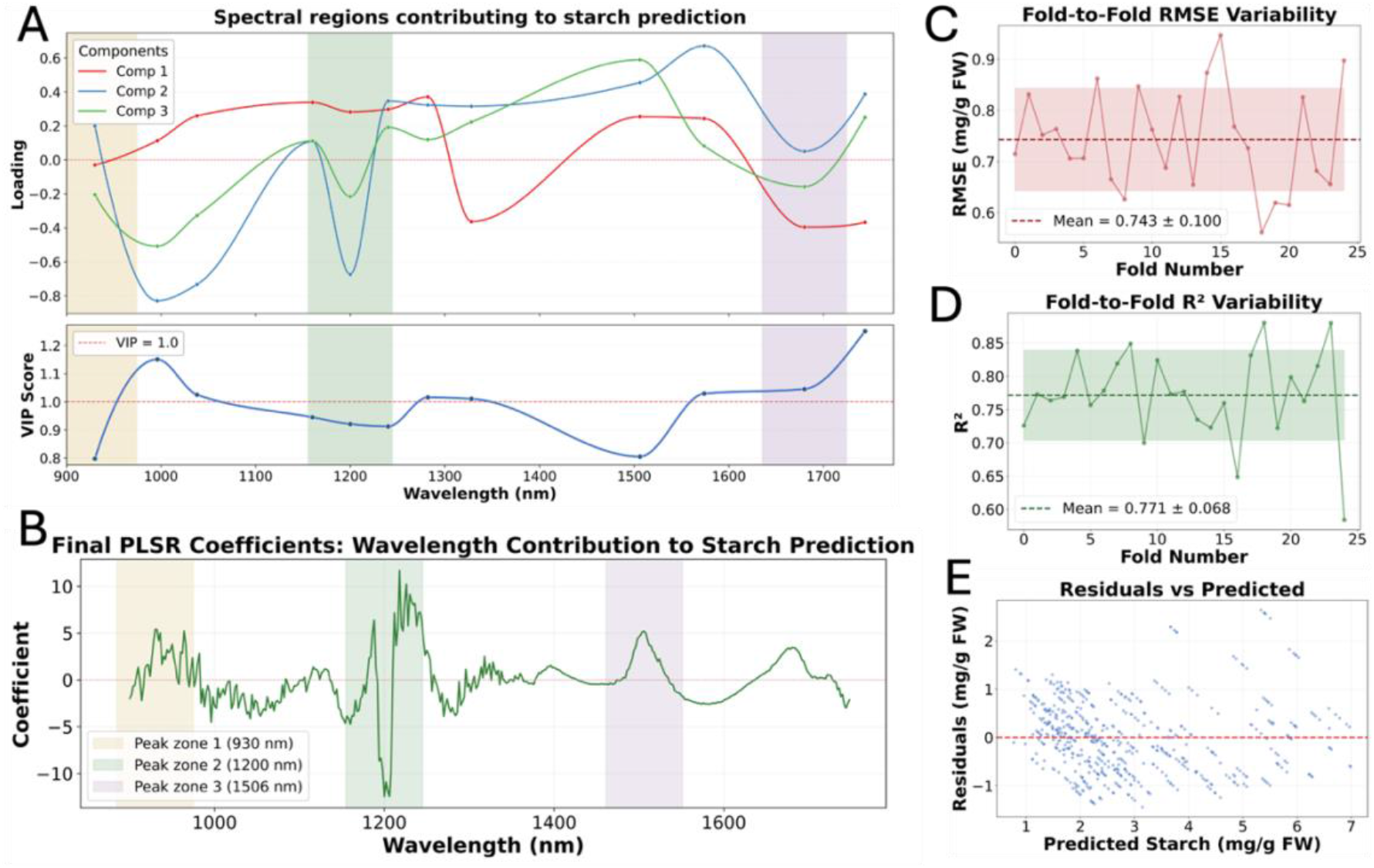
Spectral interpretability and validation diagnostics of the reduced-band PLSR model. (A) PLSR latent loading structure and Variable Importance in Projection (VIP) scores for the final constrained 12 -band model. The upper panel shows the first three latent component loadings across the selected wavelength regions, while the lower panel displays VIP scores for the corresponding wavelengths. Shaded regions indicate dominant spectral neighborhoods contributing to starch prediction. The dashed horizontal line indicates the commonly used VIP relevance threshold (VIP = 1.0). (B) Regression coefficients of the final full-spectrum PLSR model across the measured SWIR range (900–1750 nm). Positive and negative coefficient regions indicate wavelength domains contributing differentially to starch prediction. Shaded regions highlight major spectral neighborhoods associated with elevated model contribution. (C) Fold-to-fold variability in cross-validated RMSE across repeated stratified 5-fold cross-validation. The dashed horizontal line represents the mean RMSE, and the shaded region indicates ±1 SD. (D) Fold-to-fold variability in cross-validated R^2^ values across repeated stratified 5-fold cross-validation. The dashed horizontal line represents the mean R^2^, and the shaded region indicates ±1 SD. (E) Residual distribution of cross-validated predictions as a function of predicted starch concentration.

Cross-validation diagnostics demonstrated relatively stable model behavior across folds, with moderate variability in both RMSE and R^2^ values throughout repeated stratified validation runs (Fig. 3B–C). Residual analysis showed no strong evidence of systematic bias across most of the prediction range, although residual dispersion increased moderately at elevated starch concentrations (Fig. 3E).

### 3.4 Reduced-Band Optimization and Spatial Starch Reconstruction

Predictive performance improved rapidly with increasing band count before approaching a plateau at approximately 12 selected wavelengths. Beyond this point, additional spectral bands provided only marginal improvements in cross-validated R^2^ and RMSE, indicating substantial spectral redundancy within the full hyperspectral dataset (Fig. 4A). The selected 12-band configuration therefore represented the optimal tradeoff between predictive performance and spectral complexity and was selected for all downstream analyses.

**Figure 4.**
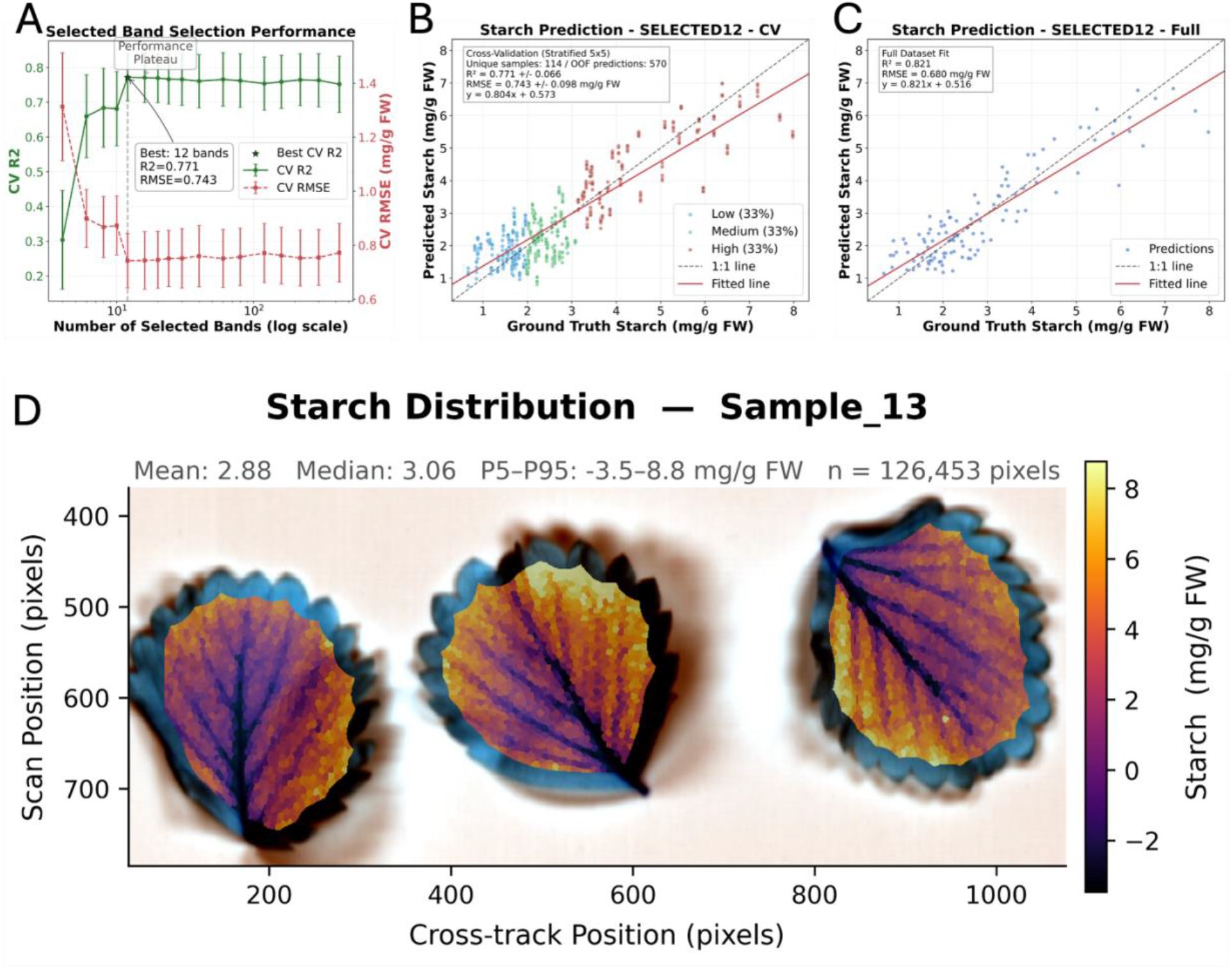
Reduced-band starch prediction performance and spatial reconstruction from SWIR hyperspectral data. (A) Cross-validated parity plot of the constrained 12-band PLSR model using repeated stratified 5-fold cross-validation (5 repeats). Points are colored according to starch concentration tertiles used for stratified sampling. The dashed black line represents the 1:1 relationship, while the solid red line indicates the fitted regression. Cross-validated predictions achieved an R^2^ of 0.771 ± 0.066 and an RMSE of 0.743 ± 0.098 mg g^−1^ FW across 570 out-of-fold predictions derived from 114 unique samples. (B) Predictive performance as a function of selected wavelength count during constrained band selection. Cross-validated R^2^ (green) and RMSE (red) are shown as mean ± standard deviation across repeated stratified cross-validation runs. Predictive performance improved rapidly with increasing band number before reaching a plateau at approximately 12 wavelengths, indicating substantial spectral redundancy within the full hyperspectral dataset. (C) Final reduced-band PLSR model fitted on the complete dataset using the selected 12 wavelengths. The fitted model was subsequently used for downstream pixel-wise starch reconstruction. The dashed black line represents the 1:1 relationship and the solid red line indicates the fitted regression. (D) Representative spatial reconstruction of starch-associated variation within intact strawberry leaflets using the final reduced-band model. Pixel-wise predictions were generated by applying the trained 12-band PLSR model to spectra extracted from segmented leaf tissue regions. Distinct intra-leaf heterogeneity was observed, including visible vein-associated spatial structure and localized variation across the lamina. Summary statistics for the reconstructed pixel distribution are shown above the map. Negative local predictions were retained to preserve transparency regarding model behavior and calibration constraints associated with bulk destructive reference measurements.

The final reduced-band PLSR model achieved a cross-validated coefficient of determination (R^2^) of 0.771 ± 0.066 with a root mean squared error (RMSE) of 0.743 ± 0.098 mg g^−1^ fresh weight across repeated stratified 5-fold cross-validation (Fig. 4B). Predicted starch concentrations showed strong agreement with destructive reference measurements across the full concentration range, although increased dispersion was observed at higher starch levels. Retraining the final model on the complete dataset yielded an overall fit of R^2^ = 0.821 and RMSE = 0.680 mg g^−1^ fresh weight (Fig. 4C), and this model was subsequently used for pixel-wise starch reconstruction.

Application of the final reduced-band model at pixel level generated spatial starch-associated maps that preserved pronounced intra-leaf heterogeneity (Fig. 4D). Distinct vein-associated structures and localized gradients across the lamina remained visible despite the substantial reduction from the original hyperspectral dataset to the final 12-band configuration. Predicted values spanned a broad local range within individual leaflets, indicating that spatially resolved spectral variation was retained following dimensionality reduction.

Localized negative predictions were observed in some low-intensity regions. These values likely reflect a structural limitation of applying a regression model calibrated on bulk destructive starch measurements to heterogeneous pixel-level spectra. Because the biochemical reference measurements integrated anatomically distinct tissue regions into single sample-level values, local pixel spectra can deviate beyond the calibration range represented by the averaged reference measurements. Negative values were therefore interpreted as comparatively low local starch-associated signal relative to the calibration range rather than physically negative starch concentrations.

## 4 Discussion

This study demonstrates that starch-associated spectral information in strawberry leaves can be captured using a selected reduced-band SWIR hyperspectral workflow while retaining sufficient predictive performance for spatial reconstruction within intact tissue. The developed pipeline combined automated hyperspectral segmentation, sample-level chemometric calibration, constrained wavelength selection, and pixel-wise inference to generate spatial starch-associated maps directly from hyperspectral image data. Importantly, predictive performance approached a plateau at approximately 12 selected wavelengths, indicating that a comparatively small subset of SWIR bands retained most of the information relevant for starch estimation.

The observed rapid performance saturation of the model with increasing band number highlights the substantial spectral redundancy present within the hyperspectral dataset. Although full hyperspectral acquisition provides dense spectral sampling, neighboring SWIR wavelengths are often highly correlated due to broad overlapping absorption structures and coupled biochemical and structural effects within plant tissue. The constrained band-selection strategy therefore sought to preserve spectral diversity while minimizing redundant neighboring wavelengths. Several selected regions aligned with known SWIR domains influenced by water absorption and carbohydrate-associated spectral structure, particularly near ∼1200 nm and within longer-wavelength SWIR regions. However, the predictive capacity of the model likely emerged from multivariate covariance patterns distributed across several spectral neighborhoods rather than isolated wavelength-specific starch absorptions alone. This is consistent with previous observations that plant biochemical estimation from SWIR spectra frequently reflects coupled optical interactions between water status, tissue structure, and carbon-related compounds rather than direct quantification of a single constituent (Burnett *et al*., 2021; Curran, 1989; Kokaly and Clark, 1999; Kokaly *et al*., 2009). A central outcome of this study was the ability to reconstruct spatial starch-associated variation within intact leaves. The generated maps revealed pronounced intra-leaf heterogeneity, including visible differences between vein-associated regions and surrounding lamina tissue. Importantly, these spatial structures were preserved despite aggressive dimensionality reduction from the original hyperspectral dataset to the final 12-band configuration. This suggests that reduced-band sensing may retain sufficient spatially resolved biochemical information for future multispectral implementations (Bai *et al*., 2024; Frey *et al*., 2020; Hou *et al*., 2025; Wang *et al*., 2021).

At the same time, the spatial predictions should be interpreted carefully. The calibration targets used for model development were derived from bulk destructive starch measurements obtained from whole-leaf samples. Consequently, the pixel-wise reconstructions do not represent direct biochemical measurements at cellular or tissue resolution. Instead, they should be interpreted as localized starch-associated spectral estimates inferred from covariance patterns learned from bulk reference measurements. This distinction is particularly important near anatomically heterogeneous regions such as veins, margins, and mixed tissue interfaces. The occurrence of negative local predictions within low-starch regions further reflects this limitation and likely arises from local spectral deviations relative to the averaged biochemical reference. Rather than artificially clipping these values, they were retained to preserve transparency regarding model behavior and calibration constraints (Burnett *et al*., 2021; Frey *et al*., 2020; Pandey *et al*., 2017).

The segmentation workflow contributed substantially to the robustness of the downstream analysis. Combining multiple spectral descriptors with voting-based thresholding enabled reliable tissue extraction across varying illumination conditions and local reflectance heterogeneity. The inclusion of mask erosion prior to spectral extraction further reduced mixed-pixel artifacts near object boundaries, which are known to influence hyperspectral regression performance. Clear bimodal separation between leaf and background pixels across all evaluated descriptor domains supported the suitability of automated thresholding without manual parameter optimization (Gao, 1996; Kruse *et al*., 1993; Liu *et al*., 2020; Otsu, 1979; Sarić *et al*., 2022).

Several limitations of the present study should be considered. First, the calibration dataset remained relatively limited in size and was generated under controlled experimental conditions using a single imaging configuration and plant system. External validation across additional cultivars, developmental stages, environmental conditions, and imaging setups will therefore be necessary to evaluate the method’s applicability and robustness of the calibration. Second, because the reference measurements integrated heterogeneous tissue regions into a single scalar value, true pixel-level validation was not possible. Future work combining hyperspectral imaging with localized biochemical sampling or histological approaches may help further resolve the relationship between local spectral structure and carbohydrate distribution. Third, the present study employed linear PLSR models, which remain widely used and robust for highly collinear spectral datasets but may not fully capture nonlinear spectral–biochemical relationships (Bai *et al*., 2024; Burnett *et al*., 2021; Mehmood *et al*., 2012; Zhang *et al*., 2023).

Despite these limitations, the results demonstrate the feasibility of reduced-band SWIR sensing for non-destructive reconstruction of starch-associated spatial variation in leaves. The identification of a compact wavelength subset retaining much of the predictive information from the full hyperspectral dataset is particularly relevant for future deployable multispectral systems, where reduced hardware complexity, faster acquisition, and lower data volumes are critical. More broadly, continuous non-destructive access to carbohydrate-associated spatial dynamics could enable new approaches for monitoring source–sink relationships, physiological state transitions, and temporal carbon allocation in controlled-environment and greenhouse production systems (Bai *et al*., 2024; Burnett *et al*., 2021; Hou *et al*., 2025; Sarić *et al*., 2022; Zhang *et al*., 2023).

## 5 Material and Methods

### 5.1 Plant Material and Experimental Design

Fragaria × ananassa cv. Favori plants (three months old) were grown in a controlled-environment vertical farming system (vGreens Grow Stack, coco coir substrate) under a 16 h / 8 h photoperiod (250 µmol m^−2^ s^−1^, 22 °C / 14 °C day/night). Three source–sink treatments were applied to generate a broad physiological range in non-structural carbohydrate (NSC) content.

Sink-limited plants (n = 16; 30 leaf samples) had reproductive organs removed and were exposed to elevated light (300 µmol m^−2^ s^−1^ for 2 days, then 400 µmol m^−2^ s^−1^ for 2 days). Source-limited plants (n = 16; 30 leaf samples) were held at 30 µmol m^−2^ s^−1^ for 4 days to deplete carbohydrate reserves. Balanced controls (n = 30; 54 leaf samples) were maintained under standard conditions (250 µmol m^−2^ s^−1^). One mature and one young leaf were collected per plant (total N = 114 leaf samples).

### 5.2 SWIR Hyperspectral Imaging

Leaf imaging was conducted across 11 time blocks spanning the full 16 h photoperiod (06:00–21:00, 1.5 h intervals) to capture diurnal variation in NSC dynamics. At six alternating blocks (blocks 1, 3, 5, 7, 9, 11: 06:00, 09:00, 12:00, 15:00, 18:00, 21:00), all three treatment groups were sampled simultaneously (five leaves per treatment, 15 leaves per block). At the five intervening blocks (blocks 2, 4, 6, 8, 10: 07:30, 10:30, 13:30, 16:30, 19:30), balanced-control plants were sampled (5 leaves per block), providing denser temporal coverage of the baseline diurnal NSC profile. This yielded a total of 114 matched imaging–biochemistry samples (source-limited: 30; sink-limited: 30; balanced: 54) (Graf and Smith, 2011; Stitt and Zeeman, 2012).

At each block, freshly harvested leaves were imaged with a push-broom SWIR hyperspectral camera (Black Industry SWIR 1.7 Pro Max, HAIP-solutions, Hannover, Germany) within 1 min of detachment to minimise post-harvest metabolic changes. Following imaging, leaves were individually weighed, snap-frozen on dry ice, and stored at −80 °C until biochemical analysis.

### 5.3 Biochemical Starch Quantification

Frozen leaf tissue was ground to a fine powder under liquid nitrogen. Approximately 100 mg homogenised tissue was extracted twice with 80% (v/v) ethanol (5 mL per extraction, 90 °C water bath, 10 min, 13,000 × g centrifugation). Supernatants were pooled for soluble sugar analysis; pellets were dried overnight for starch determination.

Starch was quantified from dried pellets using the Megazyme Total Starch Assay Kit (Rapid Total Starch protocol; K-TSTA-100A, Neogen) via sequential thermostable α-amylase and amyloglucosidase hydrolysis, followed by glucose oxidase/peroxidase (GOPOD) colorimetric detection at 510 nm (Implen™ NanoPhotometer™ NP80, Thermo Fisher Scientific, Waltham, USA) (Smith and Zeeman, 2006).

### 5.4 Image Pre-Processing and Segmentation

Hyperspectral image cubes were preprocessed prior to spectral analysis and segmentation to reduce sensor noise, suppress illumination-related artifacts, and improve spectral separability between plant tissue and background regions. All processing steps were applied to calibrated reflectance cubes. Spatial noise reduction was performed using Gaussian smoothing, followed by spectral denoising using a Savitzky–Golay filter applied along the spectral dimension (Savitzky and Golay, 1964).

To enhance spectral contrast between leaf tissue and background pixels, multiple complementary spectral descriptors were computed on a per-pixel basis. These included normalized NIR/SWIR ratios, NDWI-derived features, first- and second-order spectral derivatives, curvature-based spectral concavity metrics, SAM similarity score, and normalized deviation features in the NIR region. The descriptors were designed as leaf scores—the higher the value, the higher the probability a pixel is a leaf. The descriptors were min-max normalized to the range [0, 1]. Two masks were generated. First, for each descriptor, adaptive thresholds were determined using Otsu thresholding. Binary foreground masks generated from the individual descriptor domains were combined using a voting-based aggregation approach, in which pixels with at least one dissenting vote were classified as plant tissue. Second, the descriptors were pixel-wise averaged and the Otsu threshold of the average was computed. The pixels with the average score above the Otsu threshold were classified as foreground. The final mask was defined as an intersection of the two masks. Small isolated regions were removed by morphological filtering, and holes within connected foreground regions were filled to improve mask continuity (Gao, 1996; Kruse *et al*., 1993; Otsu, 1979).

To reduce mixed-pixel effects near object boundaries, the final binary masks were morphologically eroded by 15% of the outer fringe prior to spectral extraction. This excluded edge pixels potentially affected by background reflectance, shadowing, scattering, refraction, or partial-volume mixing. All downstream spectral averaging and pixel-wise starch prediction analyses were performed exclusively on the eroded masks.

Segmentation quality was visually inspected across all samples using pseudo-RGB projections generated from the hyperspectral cubes. Representative examples of preprocessing stages, feature maps, thresholding behavior, and final segmentation masks are shown in Fig. 1.

### 5.5 Spectral Preprocessing and Modeling

Spectral data were extracted from the eroded leaf masks and averaged per sample to obtain one representative mean spectrum per leaf. All subsequent modeling was performed using these sample-level spectra and the corresponding destructive starch measurements. Prior to regression modeling, spectra were preprocessed using Standard Normal Variate (SNV) normalization to reduce multiplicative scatter effects and baseline variation. The resulting preprocessed spectra were used as input variables for Partial Least Squares Regression (PLSR), with starch concentration as the response variable. The number of latent variables was optimized by repeated stratified 5-fold cross-validation. Samples were stratified into low, medium, and high starch concentration bins based on measured starch values to preserve the response distribution across folds. Model performance was evaluated using out-of-fold predictions, and the number of components was selected by balancing cross-validated prediction accuracy against model complexity (Barnes *et al*., 1989; Burnett *et al*., 2021; Geladi and Kowalski, 1986; Wold *et al*., 2001).

### 5.6 Constrained Band Selection

To identify a reduced spectral subset suitable for multispectral implementation, an initial full-spectrum PLSR model was trained using all available wavelength bands. Wavelengths were ranked by the absolute magnitude of their regression coefficients. A constrained greedy selection procedure was then applied to select informative wavelengths while maintaining spectral diversity. The selection procedure enforced two constraints: first, at least one wavelength had to be selected from each predefined spectral zone, and second, selected wavelengths had to be separated by a minimum spectral distance of 40 nm. The spectral zones were defined as 900–1100 nm, 1100–1450 nm, and 1450–1748 nm. For a target number of selected bands, the highest-ranked wavelength in each zone was selected first, followed by additional wavelengths chosen iteratively from the ranked list while satisfying the minimum-distance criterion. If the target number of bands could not be reached, the minimum distance threshold was relaxed stepwise in 5 nm increments (Geladi and Kowalski, 1986; Mehmood *et al*., 2012; Wold *et al*., 2001).

### 5.7 Band-Count Optimization and Model Evaluation

The constrained band-selection procedure was repeated for multiple target band counts. For each selected subset, a PLSR model was trained using only the selected wavelengths and evaluated by repeated stratified 5-fold cross-validation. Cross-validated coefficient of determination and root mean squared error were recorded for each band count (Burnett *et al*., 2021).

The final reduced-band configuration was selected based on the tradeoff between predictive performance and spectral complexity. Model performance reported in the results is based exclusively on out-of-fold cross-validation predictions. Following model validation, a final reduced-band PLSR model was fitted using the complete dataset and the selected wavelength subset. Although calibration was performed using leaf-averaged spectra paired with bulk destructive starch measurements, the resulting regression model was subsequently applied at pixel level to the hyperspectral image cubes to reconstruct spatial starch distributions within intact leaves.

For pixel-wise inference, spectra from individual pixels within the eroded leaf masks were extracted and preprocessed using the same SNV normalization pipeline applied during model training. The fitted reduced-band PLSR model was then used to predict starch-associated values for each pixel independently, generating spatially resolved starch maps (Burnett *et al*., 2021).

## 6 Acknowledgements

We thank vGreens colleagues Manuela Péres Rogriguez, Ioannis Ziogas, Entela Malkaj for their support in ground truth sampling and lab measurements. Furthermore, we thank Mark Stitt (Max Planck Institute Golm, Germany) for stimulating discussions and interpretation of results. This research was partially funded by Zentrales Innovationsprogramm Mittelstand – Durchführbarkeitsstudie (DS200615), as well as private investors of the vGreens Holding GmbH. MCG acknowledges financial support by the European Research Council under the European Union’s Horizon Europe Framework Program/ERC Advanced Grant agreement no. 101097878 (HyAngle). The authors thank Niko Mohr for discussions and feedback on the manuscript.

## 7 Author Contributions

AG performed hyperspectral image acquisition, data processing, model development, statistical analyses, and figure generation. SB conducted plant cultivation, treatment implementation, sample preparation, and destructive starch measurements and experimental design. MK contributed to spectroscopy- and optical system calibration, and data interpretation. DS developed the segmentation algorithm and contributed to image-processing methodology. JD built the hardware assembly and electrical design for the image acquisitions. HSK and BE supported with extractions methodology and result interpretation. AK advised on data analysis and interpretation of results. MCG advised on data interpretation and contributed to manuscript revision. SH conceived the study and supervised the project. All authors contributed to manuscript drafting and revisions and approved the final version.

## 9 Data and Code Availability

The datasets and code supporting the findings of this study are available from the corresponding author upon reasonable request and subject to reasonable confidentiality and intellectual property considerations.

